# An *in silico* analysis reveals eutherian NLRP3 is distinguished by conserved regulatory features absent in non-eutherian NLRP3-like proteins

**DOI:** 10.64898/2026.04.26.720791

**Authors:** Daniel M. Williams

**Affiliations:** Revvity, 8100 Cambridge Research Park, Cambridge, CB25 9TL, UK

## Abstract

NLRP3 is a cytosolic pattern recognition receptor that controls the formation of an inflammasome, a multimolecular complex that cleaves the pro-inflammatory cytokines interleukin-1β and interleukin-18 into their bioactive forms. NLRP3 has been widely assumed to be conserved across vertebrates, suggesting it plays an indispensable role within the vertebrate innate immune system. Here, gene synteny, phylogeny, and structural analysis were used to examine the evolutionary origins of NLRP3 in greater detail. This analysis revealed that ‘modern’ NLRP3, defined by gene synteny and structural features, is unique to the eutherian lineage. Non-eutherian NLRP genes share limited synteny with the eutherian NLRP3 locus and lack conservation of key features including disc forming residues, cage interfaces, and membrane binding regions. NLRP3’s characteristic regulatory architecture therefore appears to have evolved after eutherians split from marsupials 160 million years ago. Whilst further experimental studies are required to validate the conclusions drawn, these findings have potentially important implications for understanding the beneficial roles of NLRP3 signalling and suggest that eutherian specific regulatory control of NLRP3 activity may have arisen in response to distinct aspects of eutherian physiology.

## Introduction

Inflammasomes are immune signalling complexes that trigger an inflammatory response upon detection of damage associated molecular patterns (DAMPs) or pathogen associated molecular patterns (PAMPs). To date, seven receptors have been described that induce inflammasome formation, with each receptor sensing a different type of PAMP or DAMP [1]. Inflammasome receptor activation upon detection of a PAMP or DAMP initiates a cascade of events that converges on the cleavage and release of the pro-inflammatory cytokines interleukin-1β (IL1β) and interleukin-18 (IL18) [2]. Binding of the bioactive forms of these cytokines to their cognate receptors coordinates responses that serve to protect the host during the early stages of infection or tissue damage [3].

Among inflammasome receptors, NLRP3 has been the subject of intense investigation. NLRP3 belongs to the NOD-like receptor family and is defined by an N-terminal PYD domain, a central nucleotide-binding NACHT domain, and a C-terminal leucine-rich repeat (LRR) domain. It is activated by multiple structurally unrelated stimuli that, with some exceptions [4,5], induce potassium efflux [6,7], triggering conformational changes that permit interaction with the adaptor ASC and seeding of an inflammasome complex that cleaves IL-1β and IL-18 into their active forms. Aberrant NLRP3 activation contributes to the pathology of numerous disease states, including atherosclerosis [8], dementia [9] and diabetes [10]. To prevent premature activation, NLRP3 activity is subject to unusually complex regulation: a large number of post-translational modifications [11–13], protein interaction partners [14,15], and the capacity to associate with intracellular membranes [16–21] collectively keep NLRP3 activity under tight control. Structural studies have further revealed that NLRP3 transitions from inactive membrane-associated decamers, or ‘cages’ [22–25], into disk-like active assemblies [26]. These regulatory and structural features have been predominantly characterised in humans and mice over the past two decades and define what is termed here ‘modern NLRP3’.

The version of NLRP3 characterised in placental mammals has been assumed to be ancient and broadly conserved across vertebrates [27,28]. Based largely on the presence of a conserved PYD-NACHT-LRR architecture, NLRP3 orthologs have been assigned in all major vertebrate lineages, including fish, amphibians, birds, reptiles and marsupials [27], implying that NLRP3 evolved over 450 MYA when these lineages last shared a common ancestor. However, shared domain organisation alone is insufficient to classify NLRP proteins as NLRP3 orthologues. Many members of the NLRP family contain the same complement and arrangement of domains as NLRP3 yet have distinct functions in innate immunity and reproductive biology [29]. Domain architecture can therefore only be used to infer structural rather than functional orthology.

The presence of the regulatory and structural features that define modern NLRP3 provide an enhanced set of criteria to potentially identify functional NLRP3 orthologues in different vertebrate lineages. Indeed, that modern NLRP3 is broadly conserved across vertebrates is challenged by the observation that some non-mammalian NLRP3-like proteins lack the distinctive NLRP3 exon 3 region important for binding to intracellular membranes [30]. This observation raises a simple question: if these proteins lack this characteristic feature of NLRP3’s biology, are they actually NLRP3? If they are not, is NLRP3 itself or its regulatory architecture restricted to specific vertebrate lineages?

To address these questions in greater detail, this manuscript applied an integrated analysis across a broad range of vertebrate NLRP proteins to examine conserved domain organisation, gene synteny and the presence or absence of key structural features, including regions required for formation of inactive and active cage and disc complexes. These analyses reveal substantial divergence between eutherian and non-eutherian NLRP proteins and suggest that the regulatory architecture characteristic of modern NLRP3 may have emerged specifically within the eutherian lineage.

## Results

### NLRP3 resides in a unique genomic locus specific to eutherian species

Orthologous genes often share the same conserved arrangement of neighbouring genes. The presence of vertebrate NLRP genes that share synteny with human NLRP3 would support the conclusion that other vertebrate species contain NLRP3 orthologs. The NLRP3 genomic locus in humans was therefore first characterised to investigate whether vertebrate lineages contain an NLRP gene at an equivalent location. Examination of the NLRP3 locus in humans revealed that NLRP3 is commonly flanked by ZNF496 and OR2B11 (**Figure 1A**) with OR11L1, TRIM58 and OR2W3 found nearby at the end of a number of additional olfactory receptor genes (referred to as ZNF496-OR2B11-TRIM58 gene locus for short) (**Supplementary Figure 1A**). Based on currently available data, NLRP3 is present at the ZNF496-OR2B11-TRIM58 gene-block in all eutherian genomes examined. As marsupials and monotremes are the closest living relatives of eutherians, species from these lineages were checked to determine whether they contain an NLRP gene at a comparable ZNF496-OR2B11-TRIM58 locus. Although a ZNF496-OR2B11-TRIM58 gene cluster can be found in marsupials, there is no NLRP gene found between ZNF496-OR2B11 or at another location nearby (**Figure 1A**). Monotremes contain a similar gene block with TRIM58 flanked by OR2W3 and ZNF genes but also do not contain an NLRP gene within this region (**Figure 1A**). To determine whether NLRP3 is absent from this locus specifically in prototherians but present at a similar locus outside therians, representative high-quality genomes from other major vertebrate clades (birds, reptiles, amphibians) were also examined. In all cases, no comparable gene block containing NLRP3 could be found. The ZNF496–OR2B11-TRIM58 gene cluster therefore likely arose after sauropsids diverged from synapsids, with NLRP3 residing at this locus only in eutherians (**Figure 1B**).

**Figure 1.**
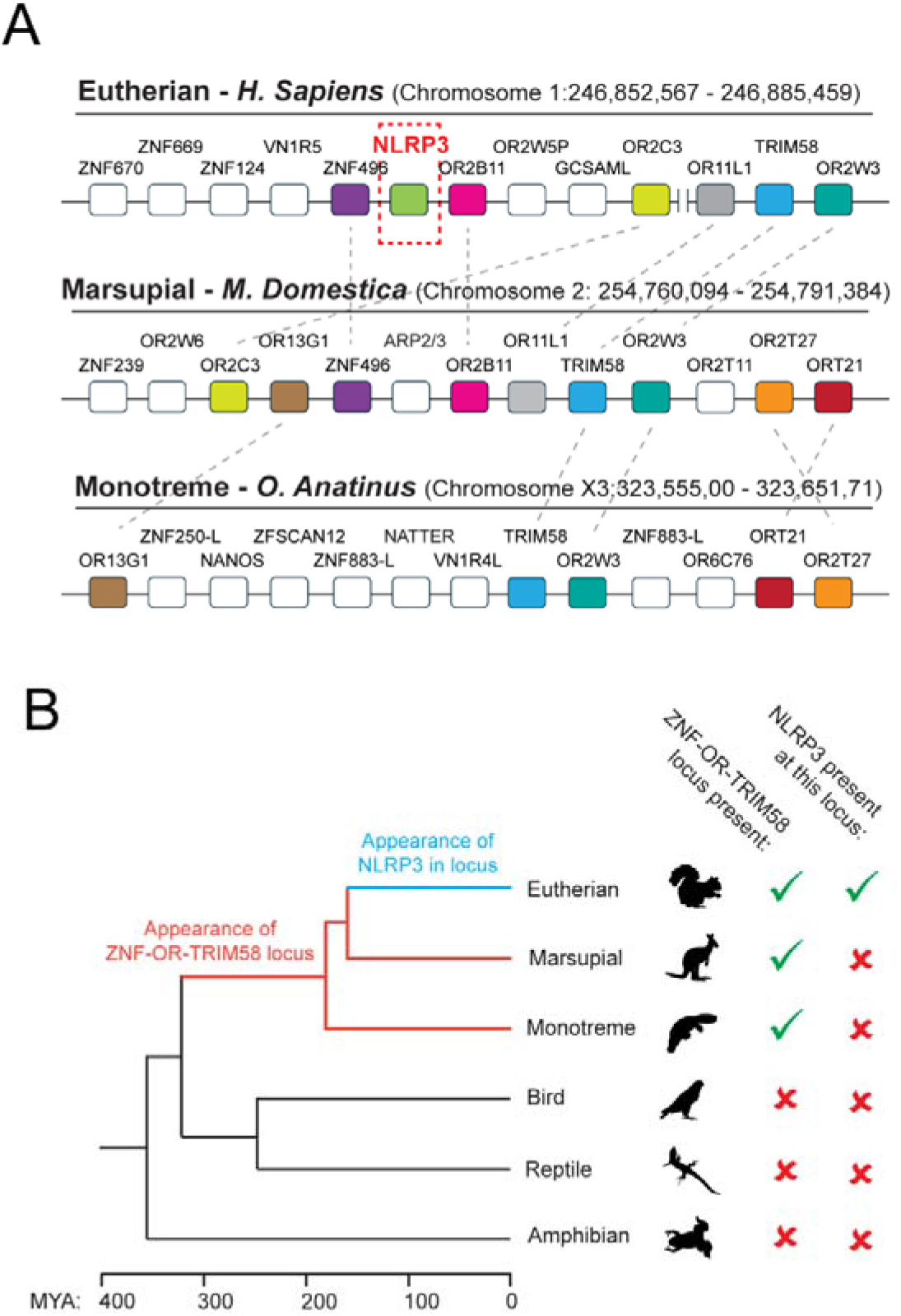
NLRP3 sits in a distinct genomic location that is unique to eutherian species. **A)** Detailed overview of the ZNF496-OR2B11-TRIM58 loci in eutherians, marsupials and monotremes. Representative chromosomal regions from *H. Sapiens* (Humans), *M. Domestica* (Grey short-tailed opossum) and *O. Anatinus* (Platypus) are displayed. Genes shared by two or more species in the same region are colour coded with homology further indicated by black dashed lines. Human NLRP3 on chromosome one is boxed and highlighted in red lettering. Genomic coordinates of each region are indicated next to species name. Additional genes between OR2C3 and OR11L1 on human chromosome one are shown in Supplementary Figure 1A. **B)** Simplified phylogenetic tree showing the proposed evolution of the ZNF496-OR2B11-TRIM58 locus that harbours NLRP3 in eutherian species. Divergence times between lineages are based on TimeTree estimates. Lineage silhouettes were obtained from phylopic.org.

### Mapping the landscape of NLRP3-like proteins in a subset of non-eutherian vertebrate lineages

To determine whether NLRP3 sits in another locus in non-eutherian vertebrate lineages, BLAST searches were performed against human NLRP3 to identify proteins showing the highest sequence similarity to NLRP3 in major vertebrate lineages (marsupials, monotremes, birds, reptiles, amphibians and fish. See **Supplementary Table 1** for full list of species analysed per NLRP protein identified). Sequence similarity to full length human NLRP3 was no higher than 60% in any of the proteins identified which was much lower than the 89% average observed between eutherian NLRP3 proteins (**Figure 2A**). Using the grey short tailed opossum as a representative marsupial genome, five NLRP-like genes could be found, consistent with a previous analysis [31]. On the basis of gene structure and synteny with other eutherian NLRP genes, three of these genes are unlikely to be NLRP3. The most likely NLRP3 candidates in marsupials are proteins referred to here as NLRP-RPS5 and NLRP-OR52I1. NLRP-RPS5 has a similar gene structure to NLRP3 (**Figure 2B-C**), containing the same intron phases and a comparable arrangement of exons encoding the PYD, NACHT and LRR domains (**Figure 2B-C**). NLRP-RPS5 is sandwiched between gene clusters consisting of UBE2M-CHMP2A-TRIM28-SLC27A5 and RNF225-RPS5-ACVRL1-ATG101 where it is also found in other marsupial species and in monotremes (**Figure 2D**), who contain an additional NLRP protein in this region. Intriguingly, the RNF225-RPS5 gene block is also present in eutherians but lacks an NLRP gene (**Supplementary Figure 1B**), consistent with either marsupials and monotremes having acquired an NLRP gene at this location or eutherians having lost one. As its naming here suggests, NLRP-OR52I1 is flanked by a number of olfactory receptors (ORs) and, similar to NLRP-RPS5, contains the same exon organisation and intron phases as eutherian NLRP3 (**Figure 2C**).

**Figure 2.**
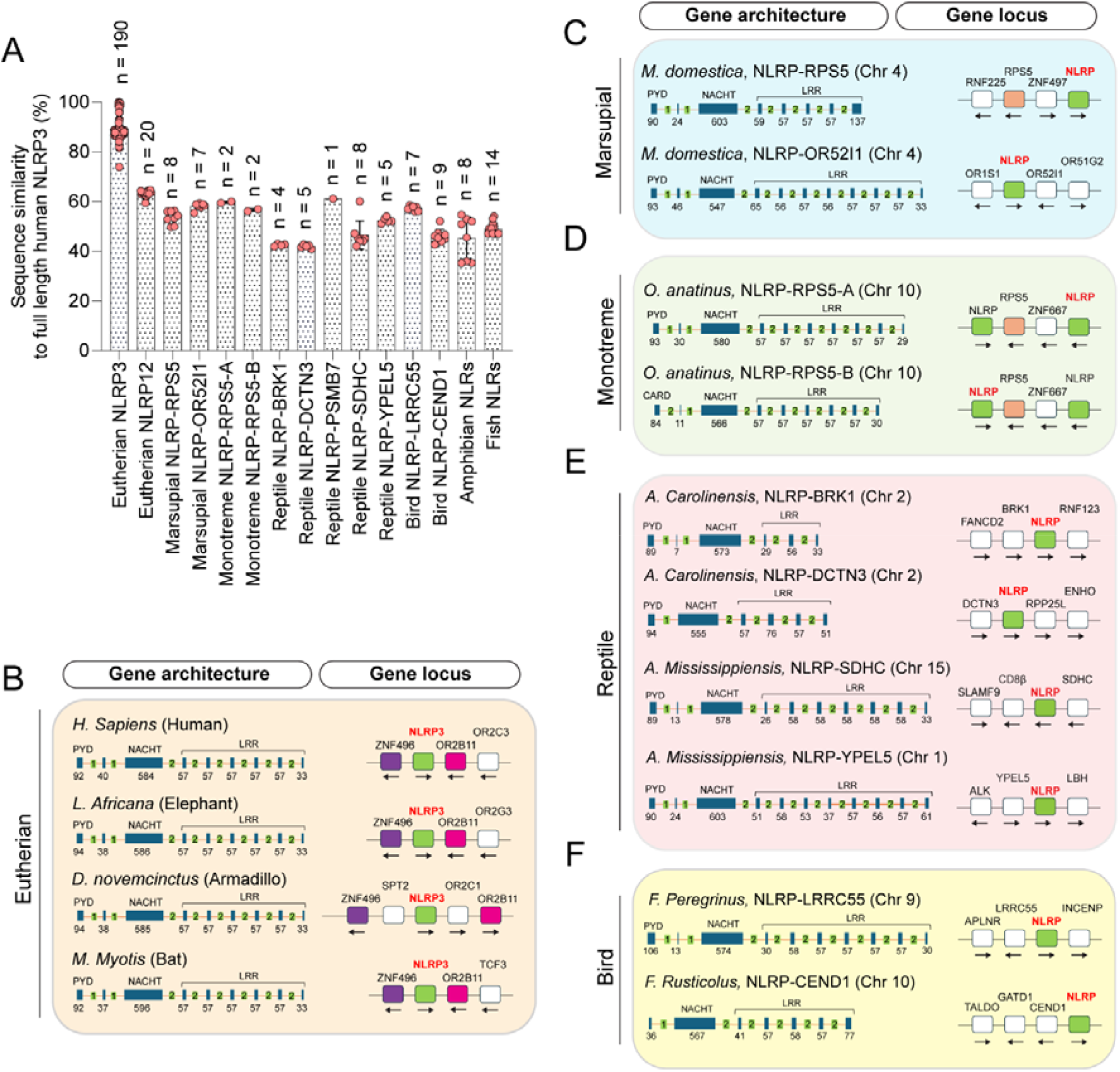
NLRP genes in non-eutherian vertebrate lineages are present at loci distinct from the eutherian NLRP3 locus. **A)** Sequence similarity of eutherian NLRP3, eutherian NLRP12 and non-eutherian NLRP proteins to the full length human NLRP3 protein. Percentage sequence similarities for amphibian and fish NLR proteins identified from BLAST searches with the highest sequence similarity to human NLRP3 are also shown. N numbers refer to the number of species analysed per protein throughout the manuscript. **B)** NLRP3 genes from representative species within the major eutherian superorders Xenarthra (Armadillo), Afrotheria (Elephant), Euarchontoglires (Human) and Laurasiatheria (Bat) showing the architecture of the NLRP3 gene (exon coding regions only) and the genes it is surrounded by in eutherian species. NLRP proteins with the highest sequence identity to human NLRP3 identified in BLAST searches against **C)** marsupials **D)** monotremes **E)** reptiles and **F)** birds. The architecture of each gene is shown with individual coding exons denoted in blue and the length of each exon in amino acids indicated underneath. Intron phases are shown in the green boxes between exons. The genomic loci of each NLRP gene are illustrated next to the architecture of each gene. NLRP proteins in non-eutherians are named by the gene they sit closest to, except for the two monotreme NLRP genes close to RPS5, which are labelled NLRP-RPS5-A and NLRP-RPS5-B. Bird NLRP-LRRC55 was previously shown to lack a PYD domain [27]. The first exon for this protein is therefore not labelled as encoding a PYD.

In other vertebrate lineages (birds, reptiles, fish, amphibians), NLRP genes with the highest sequence similarity to NLRP3 are also found at distinct genomic regions that do not share conserved synteny with the NLRP3 locus (**Figure 2E-F, Supplementary Figure 2, Supplementary Table 2**). Exceptions to this included NLRP proteins in some species of reptiles, amphibians, and fish that do have NLRP genes flanked by ZNF, TRIM or OR genes, although the assigned identities and order of the flanking genes differ markedly from the eutherian NLRP3 locus (**Supplementary Table 2, Supplementary Figure 2**). Of note, the NLRP proteins found in reptiles are not common to all reptilian orders. A gene adjacent to SDHC gene is retained in turtles, crocodiles and squamata, NLRP-YPEL is present in crocodiles and turtles, NLRP-PSMB7 is only found in crocodiles, whilst BRK1 and DCTN3 adjacent NLRP genes are only found in squamata (**Supplementary Figure 2**, **Supplementary Table 1**). These patterns of inheritance are consistent with previously documented lineage specific gains and losses of NLR genes and other inflammasome components [27,32–34].

As with the marsupial and monotreme NLRP genes identified, many of the non-eutherian NLRP genes identified in reptiles, birds, amphibians and fish had a similar gene structure to NLRP3 and shared the same intron phasing, suggestive of a common ancestral NLRP like protein, although the number of leucine rich repeats was often variable (**Figure 2E-F**). Consistent with these proteins behaving as conventional inflammasome seeding receptors, the majority of the non-eutherian NLRP proteins characterised were annotated as containing a PYD domain (**Supplementary Figure 3**) that shared moderate sequence similarity with the eutherian NLRP3 PYD (**Supplementary Figure 3B**). Exceptions to this included the N terminal domain of monotreme NLRP-RPS5-B, which was annotated as a CARD domain, and bird NLRP-LRRC55. Although specific lineages of bird that contain the LRRC55 adjacent NLRP gene do contain a PYD domain, in other bird lineages the PYD domain is replaced by a domain of unknown function [27]. NLRP proteins are therefore present in non-eutherian vertebrates, as previously documented [27].

To further resolve the evolutionary relationships between non-eutherian NLRPs and eutherian NLRP3, a phylogenetic tree was constructed using the NLRP3 NACHT domain (NLRP3^135-684^) and equivalent regions of the non-eutherian NLRPs. Eutherian NLRP3 sequences formed a strongly supported monophyletic clade with no non-eutherian NLRP proteins recovered within this clade in any replicate (**Figure 3, Supplementary Figure 4**). Bootstrap support was weaker at deeper nodes within the tree, reflecting limited phylogenetic resolution among non-eutherian proteins (**Figure 3**). Thus, although NLRP proteins are present in non-eutherian lineages, the absence of non-eutherian sequences within the NLRP3 clade would indicate that these proteins may not represent direct orthologs, consistent with residence of these genes at genomic loci that are distinct from eutherian NLRP3.

**Figure 3.**
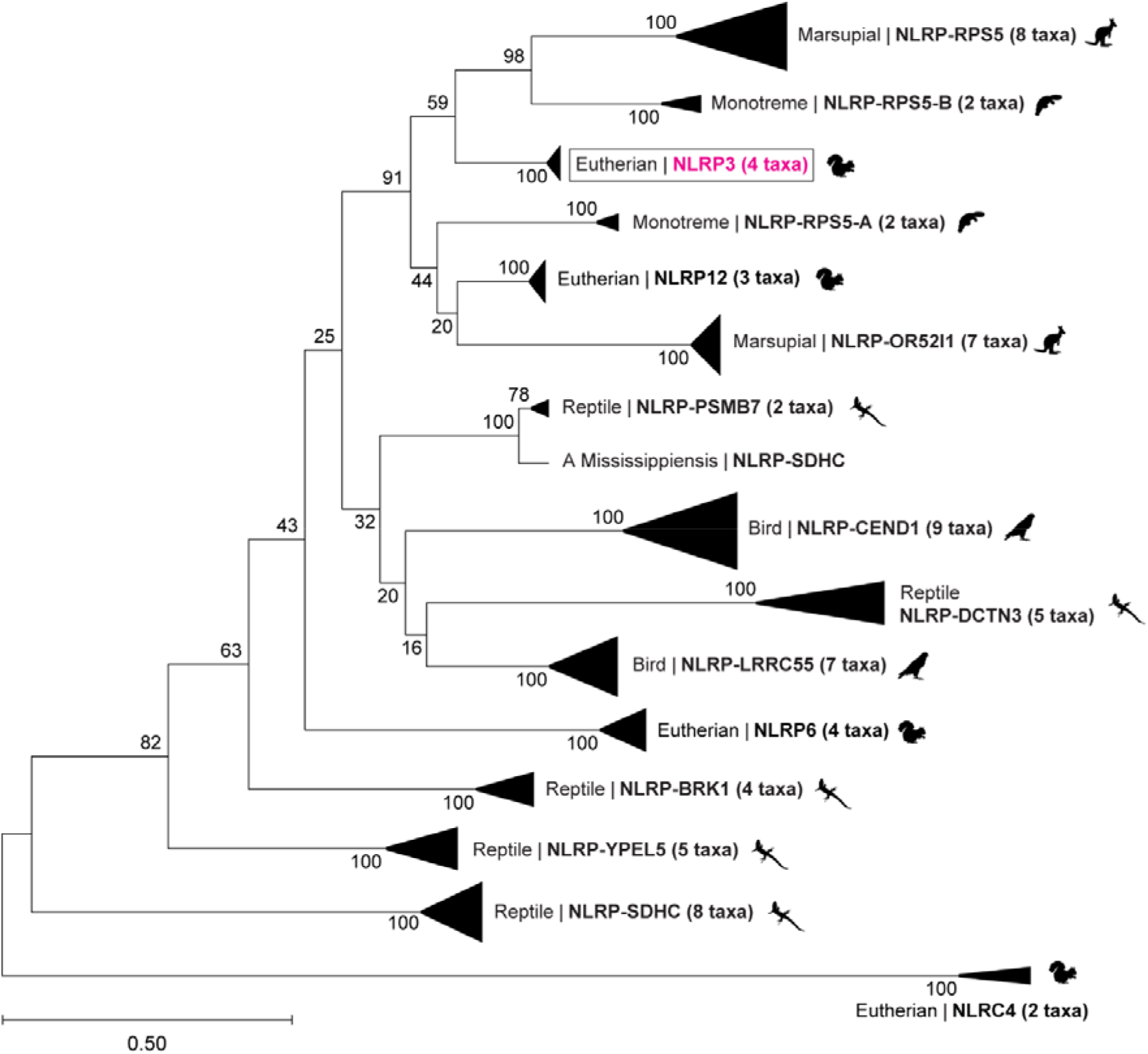
Phylogenetic analysis places non-eutherian NLRP proteins outside the eutherian NLRP3 clade. Maximum likelihood phylogenetic tree of the FISNA–NACHT region from eutherian NLRP3 proteins and non-eutherian NLRP proteins, inferred from amino acid sequences using the Jones–Taylor–Thornton (JTT) substitution model. Numbers at branch nodes indicate bootstrap support values (%) calculated from 1,000 replicates. Eutherian NLRP3 proteins form a strongly supported monophyletic clade that excludes all non-eutherian NLRP proteins. Bootstrap support is lower at deeper nodes, reflecting limited phylogenetic resolution among more divergent non-eutherian sequences. Scale bar represents 0.5 amino acid substitutions per site. The complete uncollapsed phylogenetic tree is shown in Supplementary Figure 4.

### Analysis of structural features that define eutherian NLRP3 function in non-eutherian vertebrate NLRPs

Whilst the non-eutherian NLRP proteins examined here do not share the same genetic locus as eutherian NLRP3, they may still retain the sequence and structural features that define NLRP3 function, including the FISNA and interfaces that mediate higher order assembly in its active and inactive states. Due to the wealth of structural information available [35], key features within these regions have been characterised, allowing their conservation in non-eutherian NLRPs to be examined in greater detail.

The FISNA (**<u>Fi</u>**sh-**<u>s</u>**pecific **<u>N</u>**ACHT **<u>a</u>**ssociated) domain (residues 147-220 in human NLRP3) is unique within the NLRP family to NLRP3, NLRP12 and NLRP6. For NLRP3, the FISNA is known to be tightly coupled to activation, undergoing significant structural rearrangements upon stimulation that contribute to formation of the active NLRP3 disc [26]. All non-eutherian NLRP proteins contained an identifiable FISNA with sequence similarity to human NLRP3 that was located upstream of conserved nucleotide binding motifs (Walker A, Walker B, Sensor 1, Sensor 2) (**Figure 4A**, **Figure 4C, Supplementary Figure 5A-F**). Using AlphaFold predictions of FISNA-NBD regions, structural alignments of non-eutherian NLRPs against human NLRP3 using yielded RMSD values ranging from 0.393 – 1.096 (**Supplementary Figure 6A-E**). As structural alignments were restricted to Cα atoms with AlphaFold confidence scores (pLDDT) >70, the reported RMSD values primarily reflect the NBD, for which 80–90% of residues exceeded this confidence threshold, compared with only 40–50% of residues within the FISNA domain. Nonetheless, the RMSD values were higher than structural comparisons between eutherian NLRP3s (0.18-0.28) or between NLRP3 and NLRP12 (0.373), but lower than comparisons of NLRP3 with NLRP6 (0.674) and NLRC4 (1.368). The FISNA-NBD region of many non-eutherian NLRP proteins therefore show greater structural similarity to NLRP3 than NLRP6, but less than NLRP12, placing them outside the structural range expected of direct NLRP3 orthologs.

**Figure 4.**
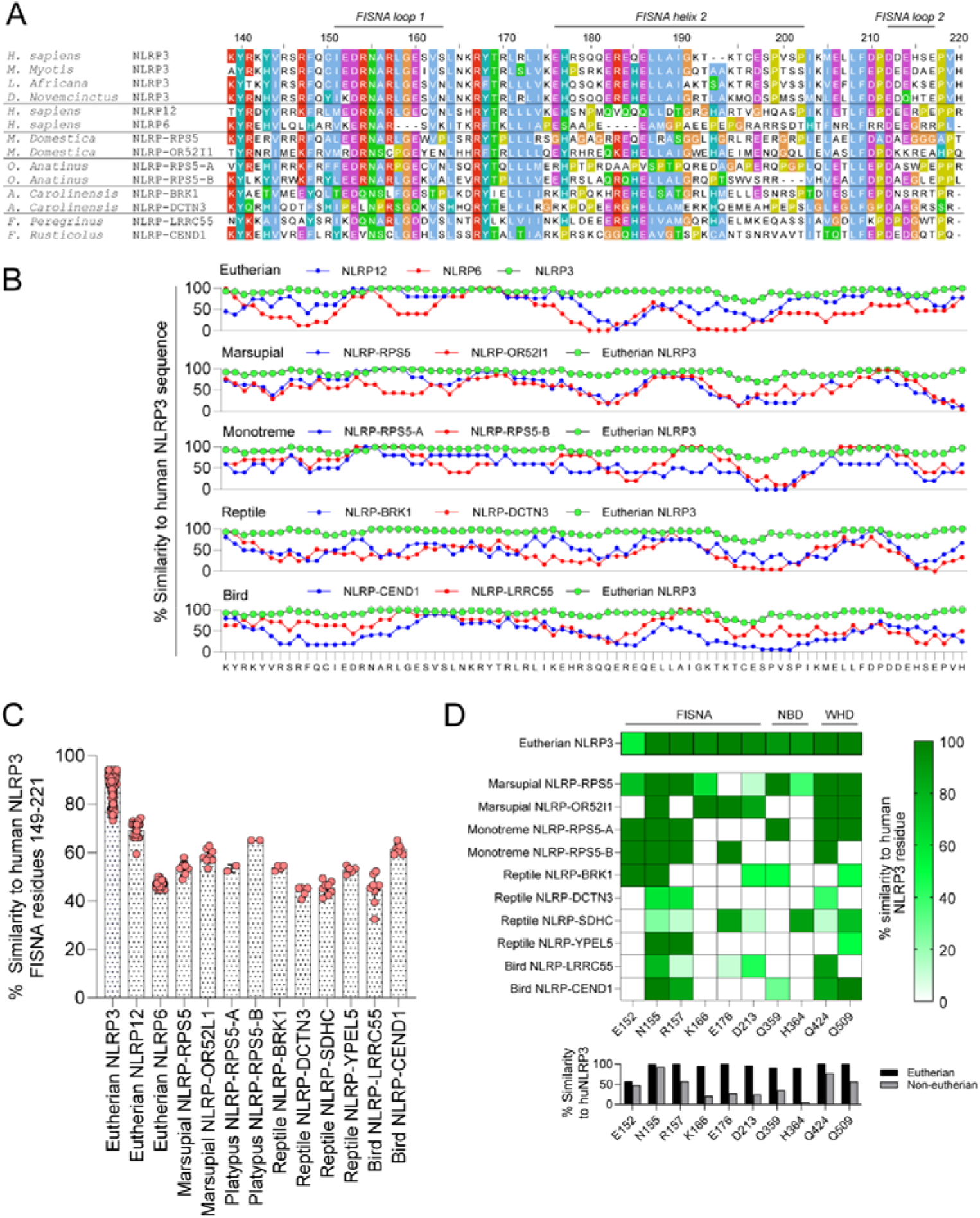
Sequence analysis of the FISNA domain and interfaces involved in formation of the active NLRP3 disc. **A)** Sequence alignments of the FISNA region from eutherian NLRP3, NLRP12, NLRP6 and the non-eutherian NLRP proteins analysed. Regions identified that undergo conformational changes in the cryo-EM structure of the active NLRP3 disk are shown above the alignment. Amino acid numbering is based on the human NLRP3 sequence. **B)** Analysis of sequence similarity of the FISNA aligned regions of non-eutherian NLRP proteins. Percentage similarity represents the average of a 5 amino acid window centred on the indicated residue. Eutherian NLRP3 similarity represents the average similarity of 190 eutherian species to the human NLRP3 sequence. Non-eutherian vertebrate lineages represent the averages from multiple species for each NLRP protein. Similarity dips in NLRP6, NLRP12 and non-eutherian NLRP proteins in regions associated with conformational changes upon NLRP3 activation, notably FISNA helix 2. **C)** Summary of percentage sequence similarity to NLRP3 of each NLRP protein across the whole FISNA region (residues 149-221 in human NLRP3). **D)** Heat map showing the percentage similarity of residues implicated in formation of the active NLRP3 disc in non-eutherian NLRP proteins.

Although an identifiable FISNA domain is present in many non-eutherian NLRP proteins, its sequence diverges from eutherian NLRP3 in key regions. Similarity is specifically reduced in the FISNA helix 2 and FISNA loop 2 regions that undergo structural rearrangement upon activation to form the NLRP3 disc (**Figure 4B**), with residues involved in disc formation also showing poor conservation in non-eutherian NLRPs (**Figure 4D**). Non-eutherian NLRPs therefore lack the residues required for disc formation as defined in eutherian NLRP3, suggesting they either do not form equivalent disc-like structures or do so through alternative interfaces.

### Physicochemical characteristics of the eutherian NLRP3 face-to-face cage-interface are not conserved in non-eutherian NLRPs

AlphaFold structural predictions of non-eutherian NLRPs were next used to examine whether interfaces important for formation of inactive NLRP3 cage complexes are conserved outside of eutherian NLRP3. Formation of the NLRP3 cage is predominantly mediated by ‘face to face’ (F2F) and ‘back to back’ (B2B) interactions between key residues within the LRR domains of opposing NLRP3 monomers (**Figure 5A**). Compared to eutherian NLRP3, the LRR domains of non-eutherian NLRP proteins are of variable length (**Figure 2A-E**). Due to alignment drifting, this makes it difficult to determine from sequence alignments alone whether residues involved in F2F and B2B interactions are conserved in non-eutherian NLRP proteins. To determine whether interfaces involved in cage formation are conserved in these proteins, the net charge and hydrophobicity of each alpha helix and beta strand within individual LRR repeats were instead calculated (**Supplementary Figure 7**). As expected, this method correctly demonstrated that the physicochemical signatures characteristic of the eutherian F2F and B2B interfaces are absent from a recently validated zebrafish NLRP3-like protein shown to be unable to form cage like structures [39] (**Supplementary Figure 8A-D**).

**Figure 5.**
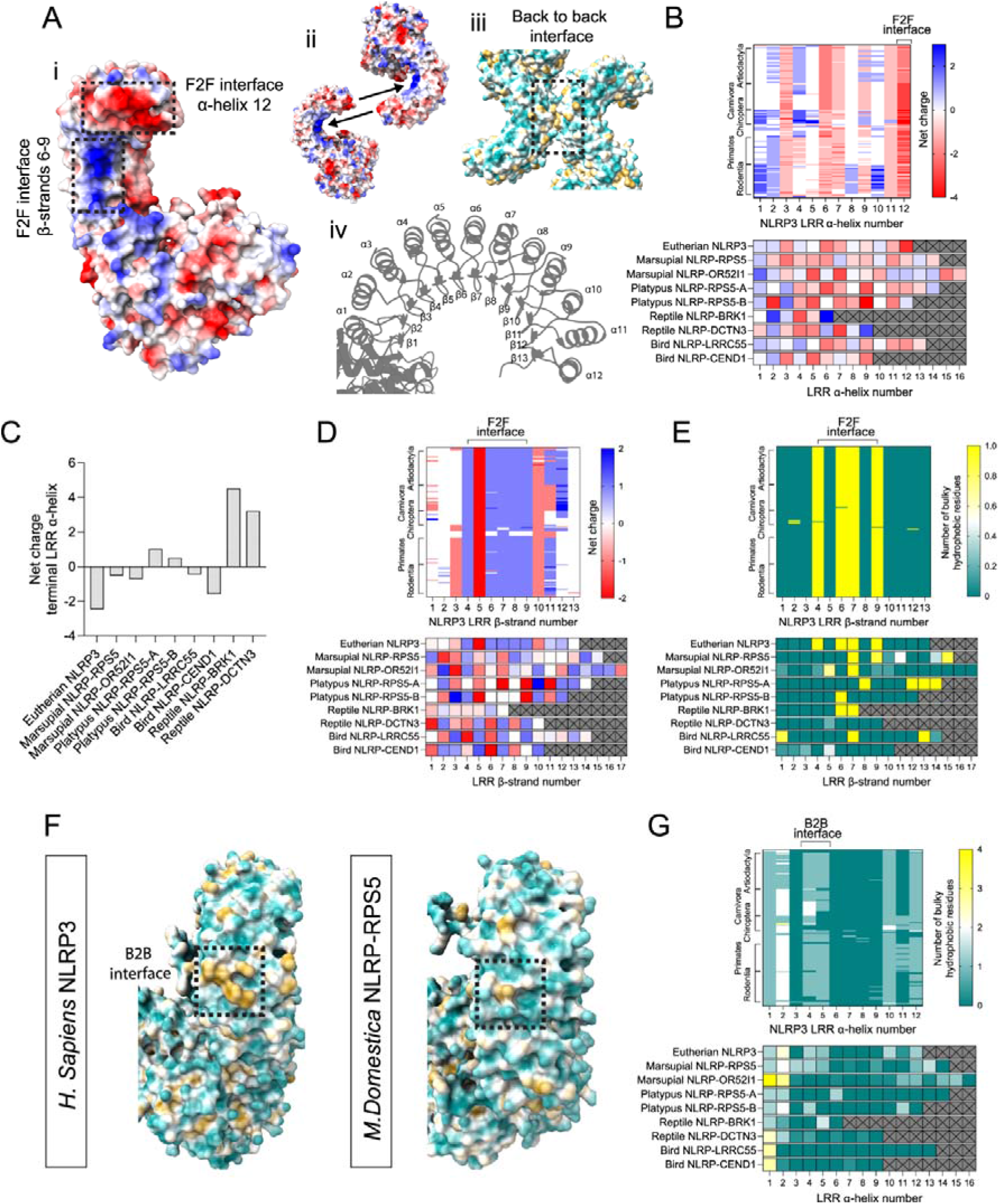
Sequence analysis of cage forming interfaces in non-eutherian NLRP proteins. **A)** ChimeraX graphic illustrating the physicochemical properties of interfaces involved in the formation of face to face and back-to-back interactions between NLRP3 monomers. Clockwise from top left: i) Electrostatic potential of the concave face of the NLRP3 LRR domain showing interfaces involved in face to face interactions between NLRP3 monomers ii) assembly logic of the face to face interface – the highly acidic terminal LRR alpha helix interacts with the concave surface of a second NLRP3 monomer iii) assembly logic of the back to back interface mediated by hydrophobic interactions iv) numbering of alpha helices and beta strands in the human NLRP3 LRR domain. Structures in (i-iii) are from (PDB ID: 7PZC), (iv) AlphaFold structural prediction of human NLRP3. **B)** Heatmap showing the net charge on convex alpha helices of the LRR domain across 190 eutherian species. Sequences are arranged by the eutherian order they belong to. The bottom panel shows a comparison of the average net charge within LRR helices calculated for each NLRP protein from non-eutherian lineages relative to eutherian NLRP3. **C)** Graph showing the net charge of the terminal LRR helix in each NLRP protein. Relative to non-eutherian NLRP proteins, the terminal NLRP3 helix is highly acidic. Heatmaps of **D)** the net charge and **E)** number of bulky hydrophobic residues (tryptophan, tyrosine, or phenylalanine) on the concave face of the eutherian NLRP3 LRR domain and non-eutherian NLRP proteins. **F)** ChimeraX graphic showing the hydrophobic patch involved in formation of back-to-back interactions in eutherian NLRP3 and its absence from *M. Domestica* NLRP-RPS5, despite the latter containing bulky hydrophobic residues in equivalent LRR alpha helices. **G)** Heat map showing the number of bulky hydrophobic residues present in convex LRR domain alpha helices of eutherian NLRP3 and non-eutherian NLRP proteins.

The F2F interface involves contacts between the terminal LRR repeat (LRR^12^) and the concave surface of the LRR, although the manner of this interaction differs between published structural models. In one model [22], the terminal LRR alpha helix (LRR^12^) contacts positively charged residues on beta-strands 5-9 of the concave LRR surface, with hydrophobic residues contributing additional contacts (**Figure 5A**). LRR^12^ in eutherian NLRP3 is highly acidic in most eutherian species (**Figure 5B-C**), facilitating the electrostatic interaction with the basic patch on beta strands 5-9, which is also highly conserved (**Figure 5D**). This patch is further enriched with bulky hydrophobic residues that again represent a conserved feature of eutherian NLRP3 (**Figure 5E**). Strikingly, non-eutherian NLRP proteins possess none of these characteristics (**Figure 5B-E, Supplementary Figure 2D, F-G**).

A second structural model places an unstructured loop rich in acidic residues (the acidic loop) at the centre of the F2F interaction [24]. However, the acidic loop also appears to be unique to eutherian NLRP3. AlphaFold structural predictions and sequence analysis revealed that although many non-eutherian NLRPs do contain an unstructured loop at an equivalent position within the LRR domain, these loops do not contain acidic character that is comparable with the loop region found in eutherian NLRP3 (**Supplementary Figure 9A-D).** Non-eutherian NLRPs therefore lack regions that correspond to the LRR F2F interfaces identified in models of the eutherian NLRP3-cage.

### Physicochemical characteristics of the eutherian NLRP3 back-to back-interface are not conserved in non-eutherian NLRPs

Like the F2F interface, the B2B interface also appears to be absent or poorly conserved in non-eutherian NLRPs. This interface centres on a pair of phenylalanines at positions 788 and 813 that are surface exposed on consecutive alpha helices (4^th^ and 5^th^ LRR helices) and form a prominent hydrophobic patch on the convex surface of the LRR domain [22–24] (**Figure 5E**). With some notable exceptions, including many species of bat (**Supplementary Figure 10A**), the arrangement of two bulky hydrophobic residues on the 4^th^ and 5^th^ LRR helices is conserved in the majority of eutherian NLRP3 proteins (**Figure 5F, Supplementary Figure 10B-C**). In contrast, hydrophobic residues positioned on consecutive convex LRR helices in a configuration compatible with the eutherian back-to-back interface are rarely observed in non-eutherian NLRPs, suggesting that these proteins may be unable to interact through an equivalent B2B interface (**Figure 5F, Supplementary Figure 2E**). The exception to this is the Marsupial NLRP-RPS5 protein that contains tryptophan and phenylalanine residues in its 4^th^ and 5^th^ LRR helices. However, AlphaFold structural predictions did not reveal a hydrophobic patch comparable to eutherian NLRP3 (**Figure 5E**), likely due to positioning of these residues within the core of the LRR fold. Together, these analyses demonstrate that the physicochemical properties underpinning both the B2B and F2F interfaces are absent from non-eutherian NLRP proteins, indicating that the inactive cage architecture may be specific to eutherian NLRP3.

### The NLRP3 membrane binding region is poorly conserved in non-eutherian NLRP proteins

Lastly, as cage formation has been shown to depend on the ability of NLRP3 to bind to intracellular membranes [22], non-eutherian NLRPs were examined to determine whether they still contain the region responsible for membrane binding despite lacking comparable cage forming interfaces. In eutherians, membrane recruitment of NLRP3 occurs through a region encoded by exon three, averaging 40 amino acids in length. This segment contains a conserved cysteine residue (Cys130 in human NLRP3) positioned between a short hydrophobic α-helix (residues 115–125) and a polybasic region (residues 131–134). Together, the α-helix and polybasic region are proposed to enable a transient membrane association that allows S-acylation of Cys-130, stabilising a pool of NLRP3 on intracellular membranes [16–20,36]. Across all eutherian NLRP3 sequences examined, the arrangement of a hydrophobic helix, central cysteine, and adjacent basic residues is preserved (**Figure 6A-B**).

**Figure 6.**
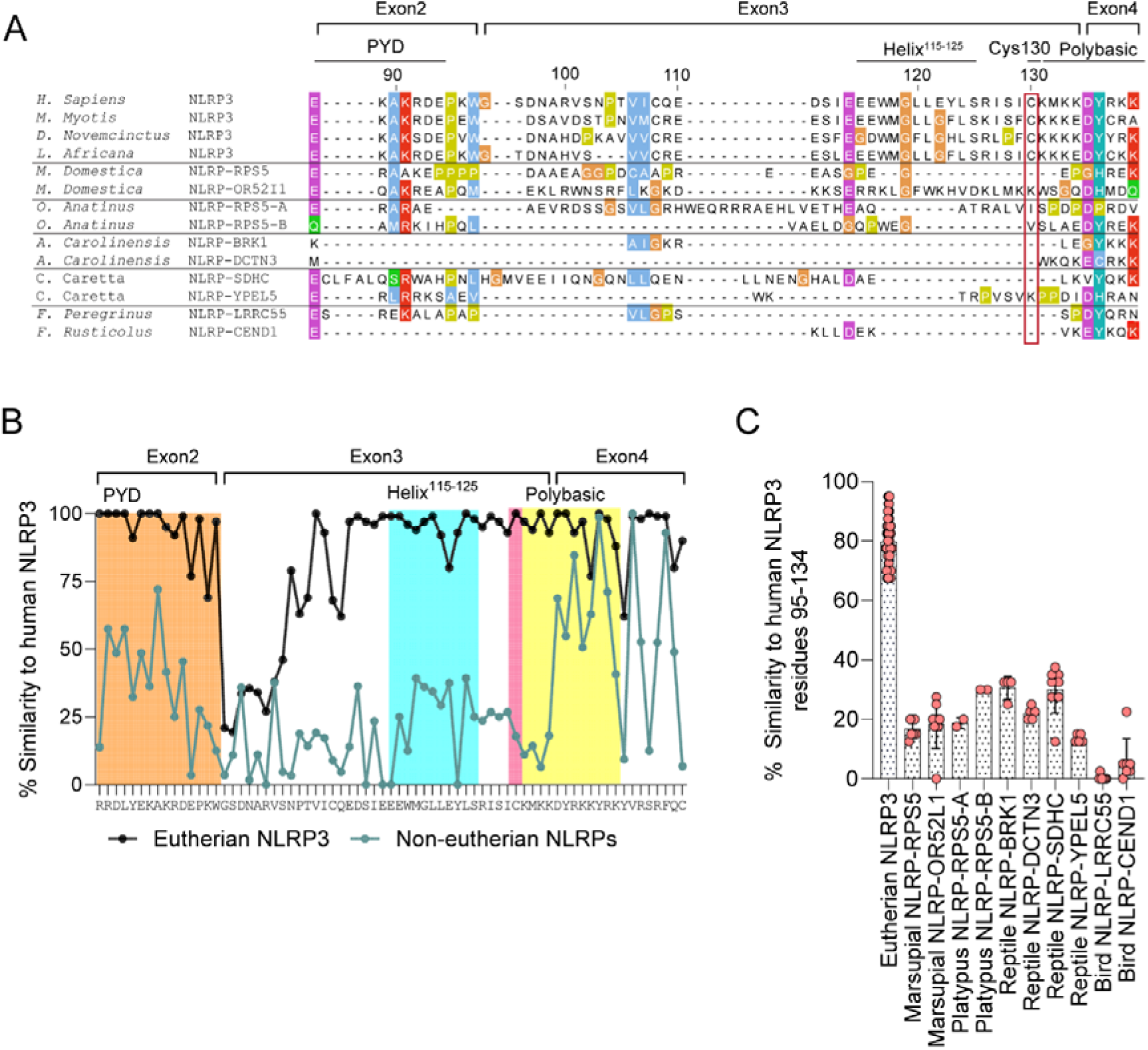
Sequence features important for association of eutherian NLRP3 with intracellular membrane are absent from non-eutherian NLRP proteins. **A)** Sequence alignments of NLRP3 and NLRP proteins from representative non-eutherian species. The position of the PYD domain, helix 115-125, Cys-130 (boxed in red) and the polybasic region are indicated based on the human NLRP3 sequence. **B)** Graph showing the % similarity of eutherian NLRP3 and non-eutherian NLRP proteins to human NLRP3 at each amino acid position of the exon3 region. Eutherian NLRP3 similarity represents the average % similarity at each position across 190 eutherian species. Non-eutherian NLRP similarity is taken from all species across marsupials, monotremes, reptiles and birds. Non-eutherian NLRP sequence similarity to NLRP3 drops off notably in the region required for NLRP3 membrane binding before increasing again at the beginning of the FISNA. **C)** Graph showing overall sequence similarity of the non-eutherian NLRP proteins to the human NLRP3^95-134^ region.

Although previous work suggested this region is absent altogether from some non-eutherian NLRP proteins [30], many of the non-eutherian NLRP proteins examined here do possess a PYD–NACHT linker encoded by a distinct exon. However, in comparison to the eutherian NLRP3 PYD-NACHT linker, these regions vary considerably in length and composition. Sequence similarity is minimal (**Figure 6C, Supplementary Figure 2H**), with none of the non-eutherian proteins analysed containing a cysteine embedded within a helix-basic motif configuration comparable with eutherian NLRP3 (**Figure 6A-B**). One exception to this is the presence of a 46 amino acid linker in marsupial OR52I1 that is rich in basic and hydrophobic residues (**Figure 6A**). Although again showing minimal sequence identity to the eutherian linker and lacking a cysteine residue that could be palmitoylated, the combination of charged and hydrophobic residues in NLRP-OR52I1 mirrors the strategy proposed to be used by NLRP3 to contact membranes prior to being palmitoylated [16–18]. The marsupial NLRP-OR52I1 protein may therefore have independently evolved a membrane binding linker region.

Collectively, these results demonstrate that the membrane-targeting region characteristic of eutherian NLRP3 is not conserved outside of this lineage. As with the B2B and F2F interfaces, the acquisition of a palmitoylation-competent membrane-binding module in the PYD-NACHT linker represents a eutherian specific specialisation and may have co-evolved in tandem with the face-to-face and back-to-back interfaces. Together, the lack of conserved synteny with the eutherian NLRP3 locus, the phylogenetic analysis, and the structural features described here, support the conclusion that several aspects of modern NLRP3s biology are unique to eutherians.

## Discussion

The evolutionary history of NLRP3 remains an understudied area yet a surprisingly common view is that NLRP3 is broadly conserved across vertebrates. Despite previous concerns regarding the use of domain architecture alone to assign orthologs [37,38], the idea that NLRP3 is broadly conserved across vertebrates is in part based on the presence of NLRP proteins containing a PYD-FISNA-NACHT-LRR domain architecture in diverse vertebrate species [27]. While NLRP proteins are present in non-eutherian vertebrates, the combined analyses conducted here indicate that these proteins do not possess the defining characteristics of the NLRP3 protein studied in humans and mice. Instead, the bona-fide ‘modern’ NLRP3 protein, defined by these same features, appears to be specific to the eutherian lineage. The results of this study, together with other recent work [39], highlight the need for more rigorous criteria in classifying NLRP proteins as genuine NLRP3 orthologues and warrant a re-evaluation of the prevailing view that NLRP3 is a pan-vertebrate inflammasome receptor.

That aspects of modern NLRP3s biology are unique to eutherians is supported by multiple lines of evidence. Consistent with another recent analysis [40], eutherian NLRP3 was found to reside in a genomic locus that no other non-eutherian vertebrate NLRP protein shares clear synteny with. The locus that harbours eutherian NLRP3 likely arose after synapsids diverged from sauropsids as a comparable ZNF496-OR2B11-TRIM58 gene cluster lacking an NLRP gene is present in marsupials and monotremes but absent from reptiles, birds, amphibians, and fish. NLRP3 was potentially acquired at the ZNF496-OR2B11-TRIM58 locus after eutherians diverged from marsupials or, alternatively, an NLRP gene may have been present at this locus in an early therian ancestor but lost from this region within the marsupial and monotreme lineages. Either way, NLRP3 has been maintained alongside ZNF496, OR2B11 and TRIM58 throughout eutherian evolution; a gene that bears the hallmarks of NLRP3 is present within the same conserved gene block in all four extant eutherian superorders.

The absence of shared synteny between NLRP3 and non-eutherian NLRP proteins would indicate that the latter proteins may not be orthologous to NLRP3. However, this interpretation is complicated by the observation that many vertebrate lineages do contain NLRP genes flanked by ZNFs, ORs or TRIMs. Although the identity, order and combination of genes present bears minimal similarity to the eutherian NLRP3 locus, marsupials, monotremes, reptiles, fish and amphibians all contain NLRP genes flanked by ZNFs, OR or TRIMs, while OR genes can also be found near bird NLRP-LRRC55. The absence of conserved synteny between eutherian NLRP3 and non-eutherian vertebrate NLRP proteins could therefore be explained by sequence divergence of ZNF, OR and TRIM genes combined with local gene rearrangements and the propensity of ZNFs, ORs and TRIMs to undergo lineage specific loss and duplication [41–45]. Given these considerations, an alternative possibility is that NLRP3 and non-eutherian NLRPs represent functionally equivalent genes that derived from an ancestral NLRP flanked by ZNF, OR and TRIMs in an early vertebrate ancestor. Rearrangement of labile ZNF, OR and TRIM rich gene clusters may have eroded any shared synteny that once existed at an ancestral locus containing a foundational NLRP3 gene.

Regardless of whether shared synteny has been obscured by subsequent genomic rearrangements, sequence and structural comparisons provide independent evidence that support the emergence of modern NLRP3 as a eutherian-specific innovation. Non-eutherian NLRP proteins lacked highly conserved regions in eutherian NLRP3 that are essential for formation of active disc structures, inactive cage complexes and binding to intracellular membranes. This would suggest that non-eutherian NLRP proteins are not subject to the same regulatory mechanisms and assembly logic that define the active and inactive eutherian NLRP3 complexes. A recent cryo-EM structure of a zebrafish NLRP3-like protein provides direct experimental support for this conclusion [39]. The zebrafish protein used interfaces that differ from the eutherian NLRP3 disc to assemble into a structurally distinct higher-order complex. In addition, this protein did not form cage structures, showed no enrichment at the Golgi and was unable to bind to NEK7 [39]. The absence of conserved regulatory features from non-eutherian vertebrate NLRP-like genes would suggest that these features did not arise in an ancestral NLRP3-like gene and instead likely evolved after eutherians diverged from marsupials. In agreement with this, NLRP3 sits in the same genomic locus and shares 80% sequence identity between Afrotheria and Xenarthra, two deeply diverged eutherian lineages that split over 100 MYA [46,47]. This would put a potential time-window for the evolution of NLRP3’s regulatory architecture between 166 to 105 MYA, after eutherians split from marsupials [48] and before the divergence of eutherians [46].

Of note, some of the interfaces required for regulatory control of eutherian NLRP3 show poor conservation in certain eutherian species. In a subset of bats, hydrophobic residues equivalent to F788 and F813 were not present, suggesting that these species may be unable to form inactive decameric cage structures. Relative to other eutherians, bats are known to have dampened NLRP3 activity via unique adaptations that include altered transcriptional priming and an NLRP3 splice variant where exon 7 within the LRR domain is skipped [49]. Alongside other bat specific modifications to innate immune signalling pathways [50,51], these alterations are in part thought to represent an evolutionary tradeoff that permits a higher tolerance of inflammatory danger signals generated by the high metabolic demands of flight [52,53]. The absence of cage forming residues described here may represent a similar evolutionary strategy some species of bats have employed to limit, but not completely abolish, NLRP3 activity.

A key unresolved question from this study is whether non-eutherian NLRP proteins are genuinely capable of responding to stimuli that induce potassium efflux and thus whether these proteins are functionally equivalent to eutherian NLRP3. An inability of the NLRP proteins identified here to respond to canonical NLRP3 stimuli would suggest that it is only eutherians that have evolved a potassium sensitive inflammasome receptor. However, despite recent progress [54], how NLRP3 senses potassium depletion is not fully understood. At present it is therefore difficult to determine from sequence and structural features alone whether non-eutherian NLRP proteins can respond to canonical NLRP3 stimuli. The sequence features used here to define NLRP3 appear to be additional regulatory nodes fixed onto NLRP3 to modulate the probability of activation in response to stimulation. For instance, human NLRP3 can activate in the absence of membrane binding and the formation of higher order inactive cage complexes, albeit at a slower rate [55]. Thus, it is possible that the NLRP proteins present in non-eutherians retain the capacity to sense ionic stress, which is consistent with reports of fish macrophages responding to some canonical NLRP3 stimuli [38,56]. Eutherian NLRP3 and non-eutherian NLRPs could represent derivatives of an ancestral potassium sensitive NLRP protein that evolved in an early vertebrate ancestor, with the distinctive regulatory features of modern NLRP3 characterised over the past two decades built on top of this ancestral base model.

What aspects of eutherian physiology might have necessitated the evolution of additional regulatory controls such as cage formation and membrane binding? Despite the problems that inappropriate NLRP3 activation can cause, the presence of a highly conserved NLRP3 protein at the ZNF496-OR2B11 locus across eutherian species suggests that retention of modern NLRP3 and its distinct regulatory mechanisms provides a clear benefit. Although it is likely that non-eutherian NLRP proteins have developed their own regulatory mechanisms, the presence of controls specific to eutherian NLRP3 raises the possibility that more flexible regulation of inflammasome activation became necessary in eutherians. Since the closest living relatives of eutherians, marsupials, do not encode a protein with the defining features of modern NLRP3, an interesting perspective on the above question is to consider the physiological traits that differ between these lineages. Examples of eutherian specific traits include adaptations associated with gestation [57] and the emergence of UCP1-dependent brown adipose tissue thermogenesis [58].

One speculative explanation for the emergence of the distinctive regulatory features of eutherian NLRP3 is that the evolution of extended gestation and placental biology imposed new pressures on inflammatory signalling that shaped the evolution of eutherian NLRP3. Eutherian pregnancy involves co-ordinated pro- and anti-inflammatory phases [57]. Successful implantation of eutherian embryos is dependent on an inflammatory reaction that allows attachment to the uterine epithelium [57]. To prevent rejection of the embryo by the mother, implantation is followed by a prolonged period of immune tolerance before returning to a pro-inflammatory state to facilitate parturition [59,60]. In contrast to eutherians, a prolonged anti-inflammatory phase is not required in marsupials as the embryo only briefly attaches to the uterine wall prior to parturition [57,60]. The anti-inflammatory state is therefore exclusive to eutherians but presents a unique challenge: the immune system must be corralled to tolerate the developing foetus whilst retaining its ability to protect the mother against infection [61]. To balance these needs, the eutherian immune system is known to have undergone adaptations that include the rewiring of immune regulatory networks and the acquisition of eutherian specific genes [62–64]. For eutherian NLRP3, structural features such as membrane binding and cage formation may have increased the dynamic range of NLRP3 signalling, permitting prolonged restraint of inflammasome activity during immune tolerance whilst retaining the capacity for rapid and robust activation in response to severe infection or tissue damage that threaten maternal or foetal survival.

Overall, whilst the findings presented in this paper require further experimental evidence to fully validate the conclusions drawn, the results strongly indicate that modern NLRP3, as defined by its regulatory architecture and genomic context, is a relatively recent evolutionary innovation restricted to eutherian mammals.

## Methods

### Analysis of eutherian and non-eutherian NLRP sequences

For eutherian NLRP3 proteins, NLRP3 sequences were downloaded from the NCBI orthologs database in July 2024. In total, NLRP3 sequences from 190 eutherian NLRP3 species were used for the analysis. To identify non-eutherian NLRPs with the highest sequence similarity to human NLRP3, the BLASTp server was used. Human NLRP3 (Q96P20) was used to search the marsupial, monotreme, reptile, bird, amphibian, and fish lineages. Two or more hits with the highest sequence similarity and a minimum of 85% sequence coverage were selected for further analysis. For NLRP6 and NLRP12 sequences, representative sequences were identified by BLAST searches of human NLRP6 or NLRP12 against the four major eutherian superorders. A full list of accession numbers for each eutherian and non-eutherian NLRP protein analysed in this study is provided in **Supplementary Table 1**. Sequence alignments were performed in Jalview [65] using MAFFT [66] and are provided in **Supplementary Table 3**. Intron / exon organisation and gene synteny were visualised for each gene using the NCBI genome viewer. Where possible, gene synteny for NLRP genes was checked using Genomicus. Intron phases were annotated manually using exon files for NLRP gene transcripts downloaded from the genome viewer on NCBI.

### Phylogenetic tree construction

Sequences used for the phylogenetic tree were first aligned in Jalview. The FISNA-NACHT region from human NLRP3 was selected based on the human NLRP3 FISNA-NACHT (residues 147-650), with the PYD and LRR domains removed manually in Jalview. Any large gaps across sequences were removed using Gblocks (phylogeny.fr). The resulting sequences were downloaded and imported into MEGA for phylogenetic analysis. The protein phylogenetic tree was constructed in MEGA using the maximum likelihood method with 1000 bootstrap replicates and the Jones-Taylor-Thornton substitution model.

### Structural models and alignments

Structural models for eutherian NLRP3 and representative non-eutherian NLRP proteins were generated using AlphaFold3 (alphafoldserver.com). Structures were downloaded and visualised in PyMol [67] and ChimeraX [68]. Structural comparisons and RMSD values for FISNA-NBD domain alignments were performed in PyMol. The FISNA-NBD was identified manually through visual inspection of each NLRP protein. The NBD was defined from the first to the last β-strand of the Rossmann-like fold, ending before the transition into the HD1 α-helical region. The ‘super’ function was used in PyMol to align two FISNA-NBD structures, centred on Cα atoms of each amino acid. Amino acid residues and regions with low structural confidence (pIDDT value < 70) were excluded from alignments.

### LRR domain analysis

For each representative non-eutherian NLRP protein or human NLRP3, the position of individual beta-strands and alpha helices within the LRR domain were identified using the ‘sequence’ feature in ChimeraX, which highlights α-helices and β-strands throughout the protein. Each structure was inspected visually to confirm the presence of an α-helix or β-strand at the indicated position. The position of each LRR α-helix or β-strand identified was mapped onto the respective NLRP amino acid sequence in Microsoft Excel. This was used as a reference sequence to infer the position of LRR β-strands and α-helices for orthologous NLRP proteins from other species within each lineage. For example, the position of α-helices or β-strands within the AlphaFold predicted LRR domain for *M. Domestica* NLRP-RPS5 was used to determine α-helix and β-strand positions in NLRP-RPS5 proteins from other marsupial species. Per 28/29 amino acid LRR repeat, sequences were checked to ensure consistent alignment with the canonical LRR repeat sequence identified in NLRP3 as a guide (e.g. ^1^**L**xx**L**x**L**xxxx**L**xxxxxxx**L**xxx**L**xxxxx^28^). The net charge of the β-strand or α-helix were calculated by subtracting the number of basic residues from the number of acidic residues present within each defined helix or strand. The number of bulky hydrophobic residues were calculated by summing the number of phenylalanine, tryptophan or tyrosine residues present within each defined region. An average net charge and number of bulky hydrophobic residues within each helix or beta strand was calculated for all species included for each NLRP protein and plotted as a heatmap.

## Supporting information

Supplementary Table 1. Sequence identifiers for the eutherian NLRP3 and non-eutherian NLRP proteins analysed.

Supplementary Table 2. Sequence identifiers for amphibian and fish NLR proteins.

Supplementary Table 3. Sequence alignments of eutherian NLRP3, marsupial, monotreme, reptile, bird NLRPs and eutherian NLRP12.

AlphaFold structures

Reviewer reports

## Data availability

All AlphaFold structural models and sequences used for analysis are provided as supplementary files. All other data is available upon request to the corresponding author.

## Acknowledgements

I am grateful to Professor Andrew Peden (University of Sheffield) for comments and feedback on the manuscript.

## Supplementary Figures

**Supplementary Figure 1.**
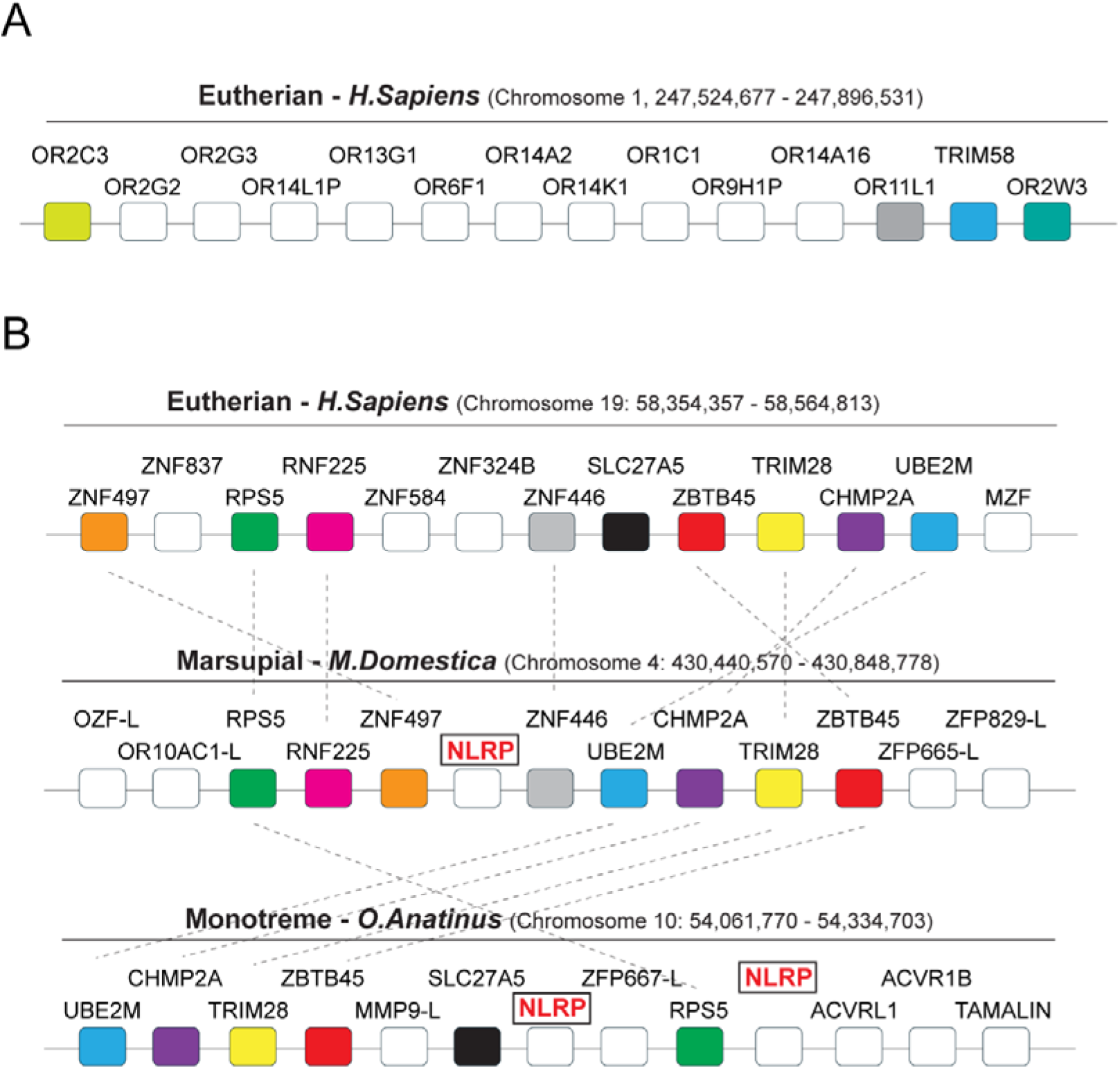
Comparative genomic analysis of regions harbouring NLRP genes in mammalian lineages. **A)** Extended view of the region on human chromosome one between OR2C3 and OR11L1. Several olfactory receptor gene duplications are present close to the human NLRP3 locus. **B)** Schematic of a homologous chromosomal region present in eutherians, marsupials and monotremes defined by the presence of RPS5 and a ZBTB45-TRIM28-CHMP2A-UBE2M gene block. This region contains either one or two NLRP genes in marsupials and monotremes, respectively. Eutherians lack an NLRP gene in this region. Orthologous genes between species are colour coded with homology between species indicated by dashed black lines.

**Supplementary Figure 2.**
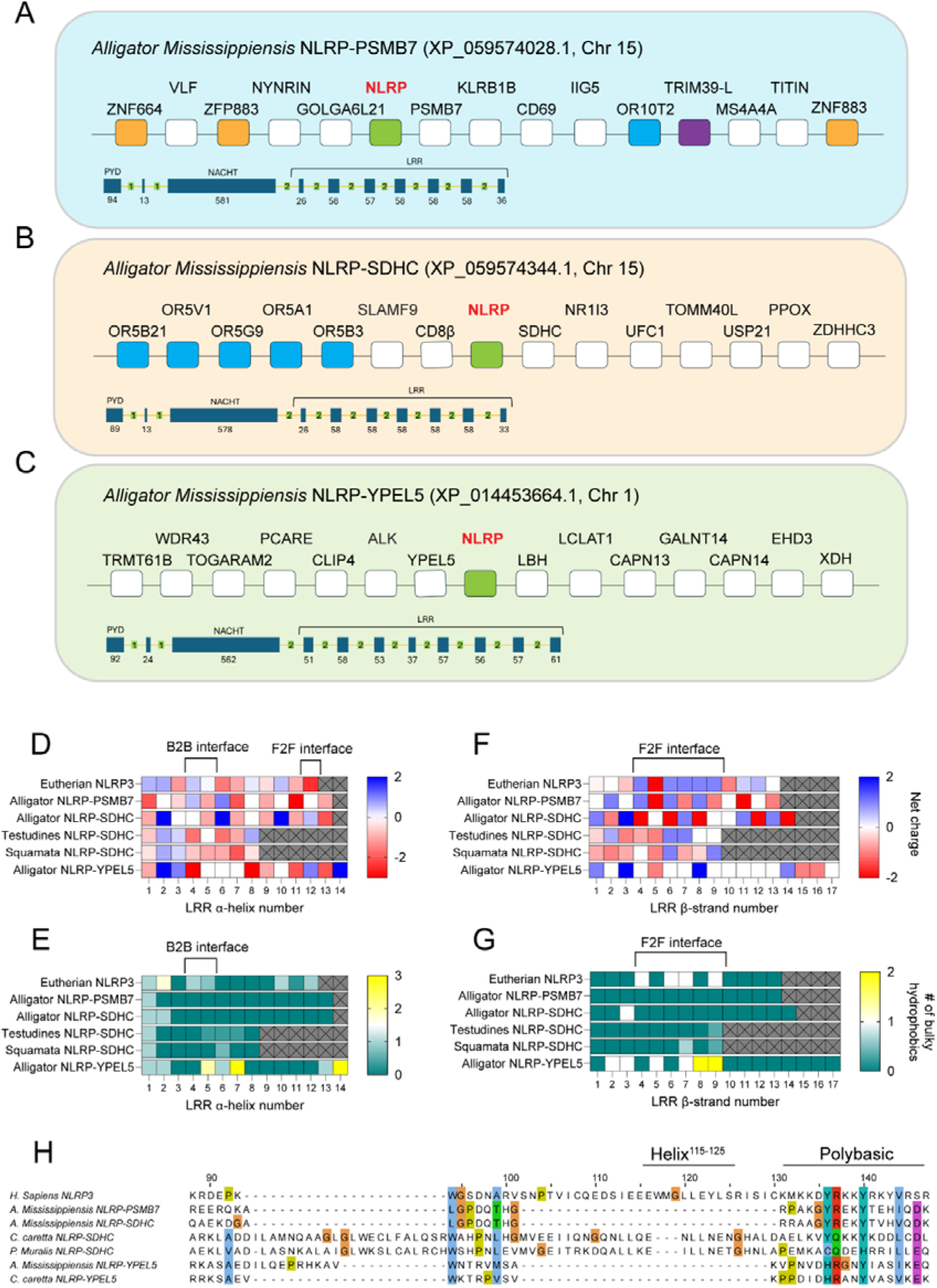
Analysis of additional reptilian NLRP proteins. Expanded view of the genomic location for the reptilian NLRP proteins **A)** NLRP-PSMB7 **B)** NLRP-SDHC and **C)** NLRP-YPEL5. *Alligator Mississippiensis* is used as a representative example. These genes show intermittent patterns of inheritance across reptilian orders (see also Supplementary Table 1). **D)** Net charge and **E)** number of bulky hydrophobic residues within each alpha helix of the indicated LRR domain. **F)** Net charge and **G)** number of bulky hydrophobic residues within each beta strand of the indicated LRR domain. **H)** Sequence alignments of the human NLRP3 membrane binding region with the indicated reptilian NLRP protein. Residue numbering is based on the human NLRP3 sequence.

**Supplementary Figure 3.**
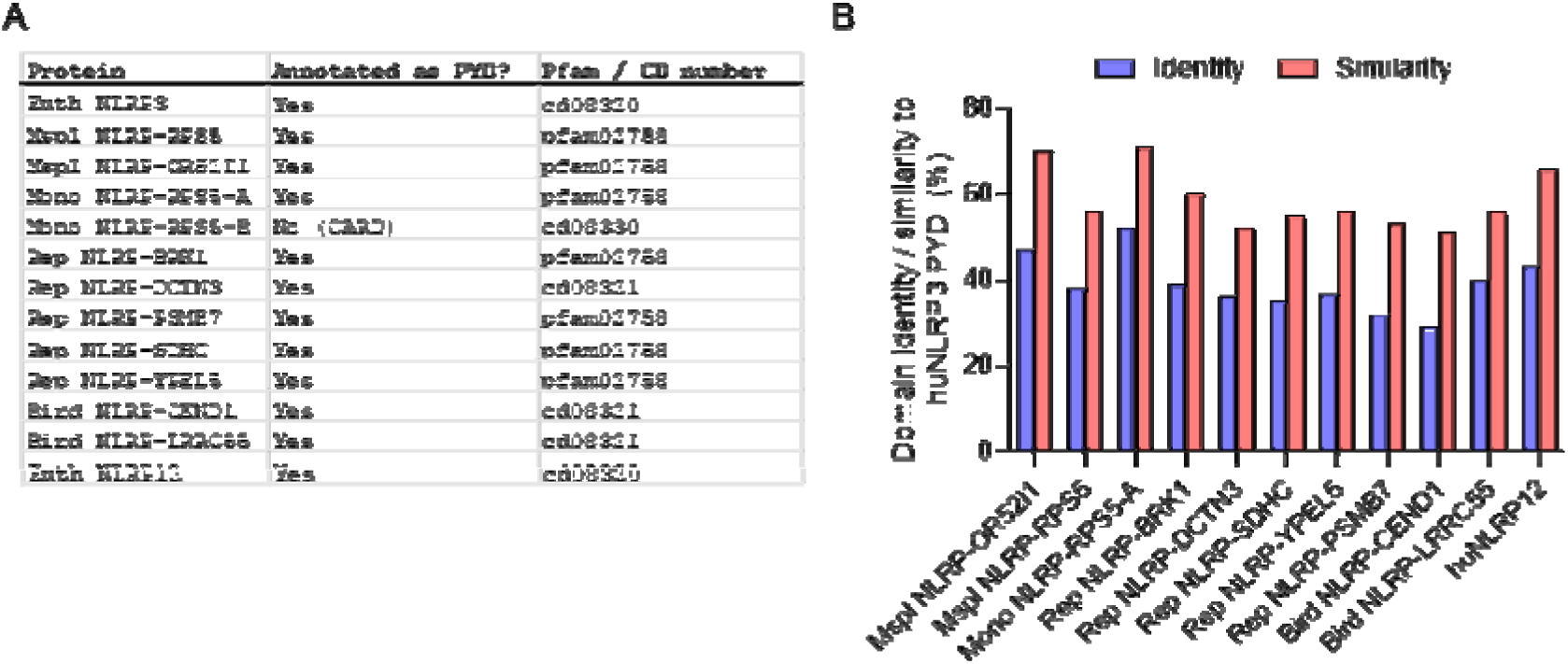
Analysis of the N terminal effector domains of non-eutherian NLRP proteins. **A)** Domain annotations for the N terminal effector domain of each non-eutherian NLRP protein and assigned PFAM or CD numbers. **B)** Sequence identity and similarity of non-eutherian NLRP PYD domains to the human NLRP3 PYD domain. Sequence alignments were calculated from one representative species for each protein. No significant similarity to the huNLRP3 PYD was found for the N terminal effector domain of monotreme NLRP-RPS5-B.

**Supplementary Figure 4.**
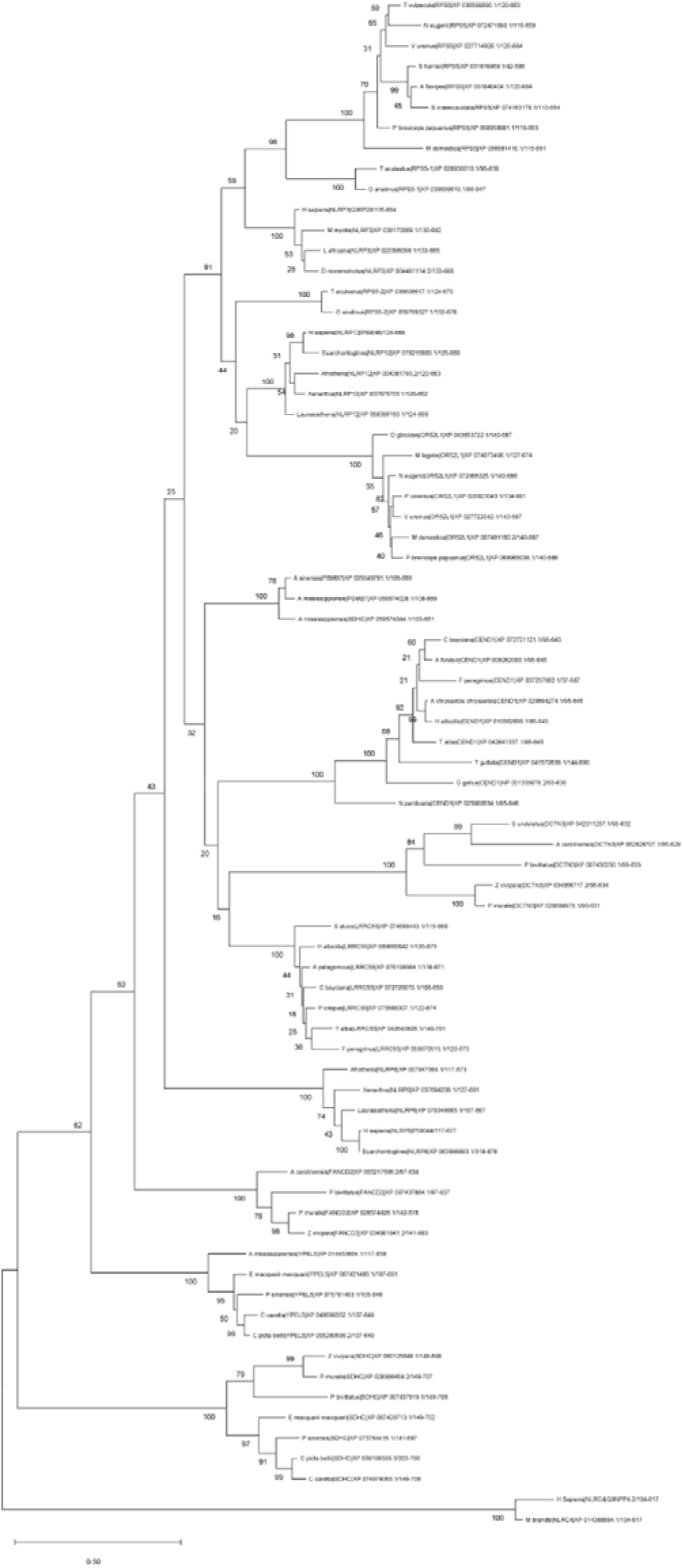
Uncollapsed phylogenetic tree of the FISNA-NACHT domain from eutherian NLRP3, NLRP6 and NLRP12 and non eutherian NLRPs. The tree was constructed using the maximum likelihood method with 1000 bootstrap replicates. The percentage of trees in which the indicated sequences clustered together is shown adjacent to each branch. Related to Figure 3.

**Supplementary Figure 5.**
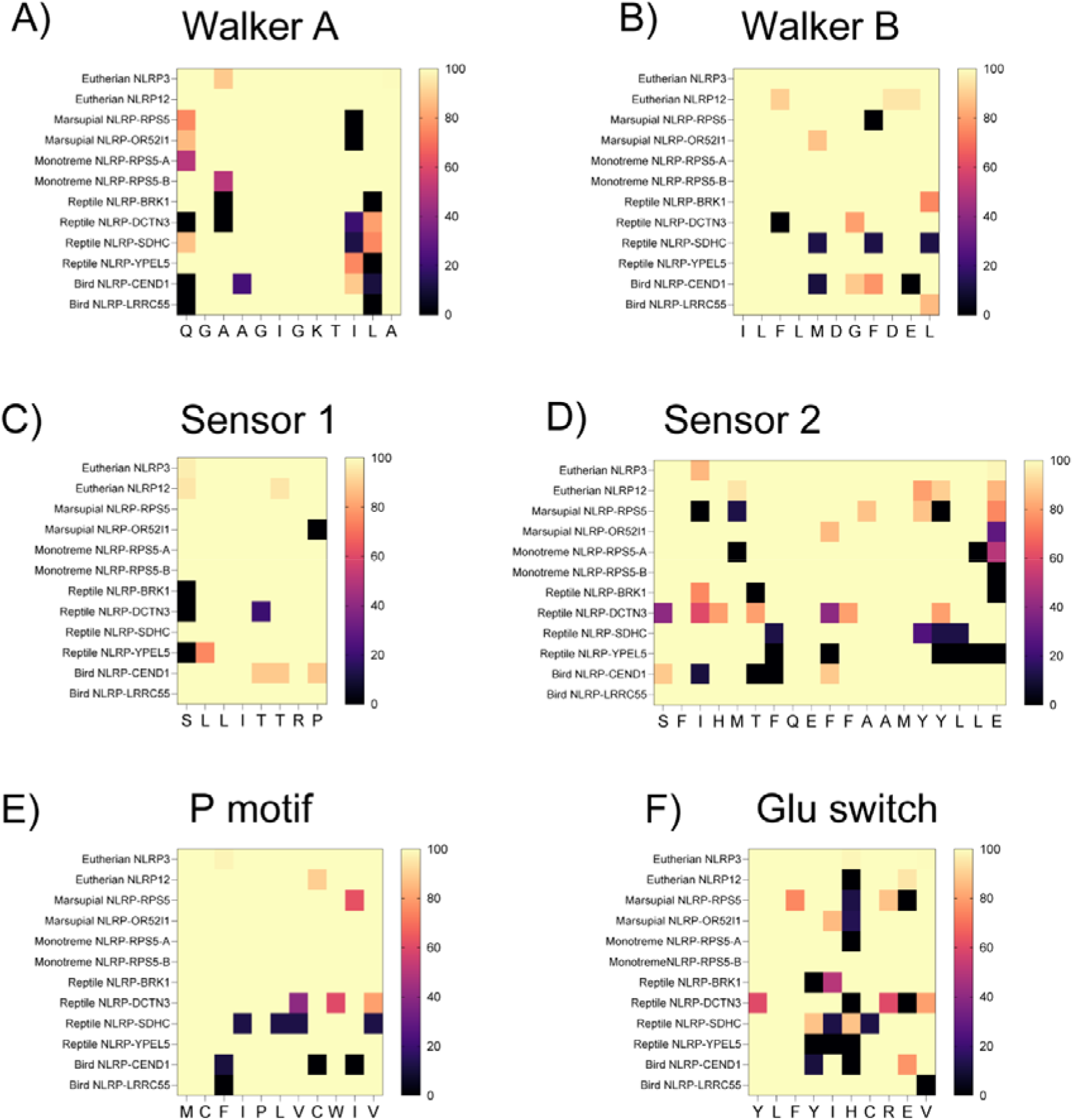
Conservation of nucleotide binding motifs in eutherian NLRP3 and non-eutherian NLRPs. Heatmaps for non-eutherian NLRPs showing the percentage of species with a similar residue to the human NLRP3 sequence at the indicated position.

**Supplementary Figure 6.**
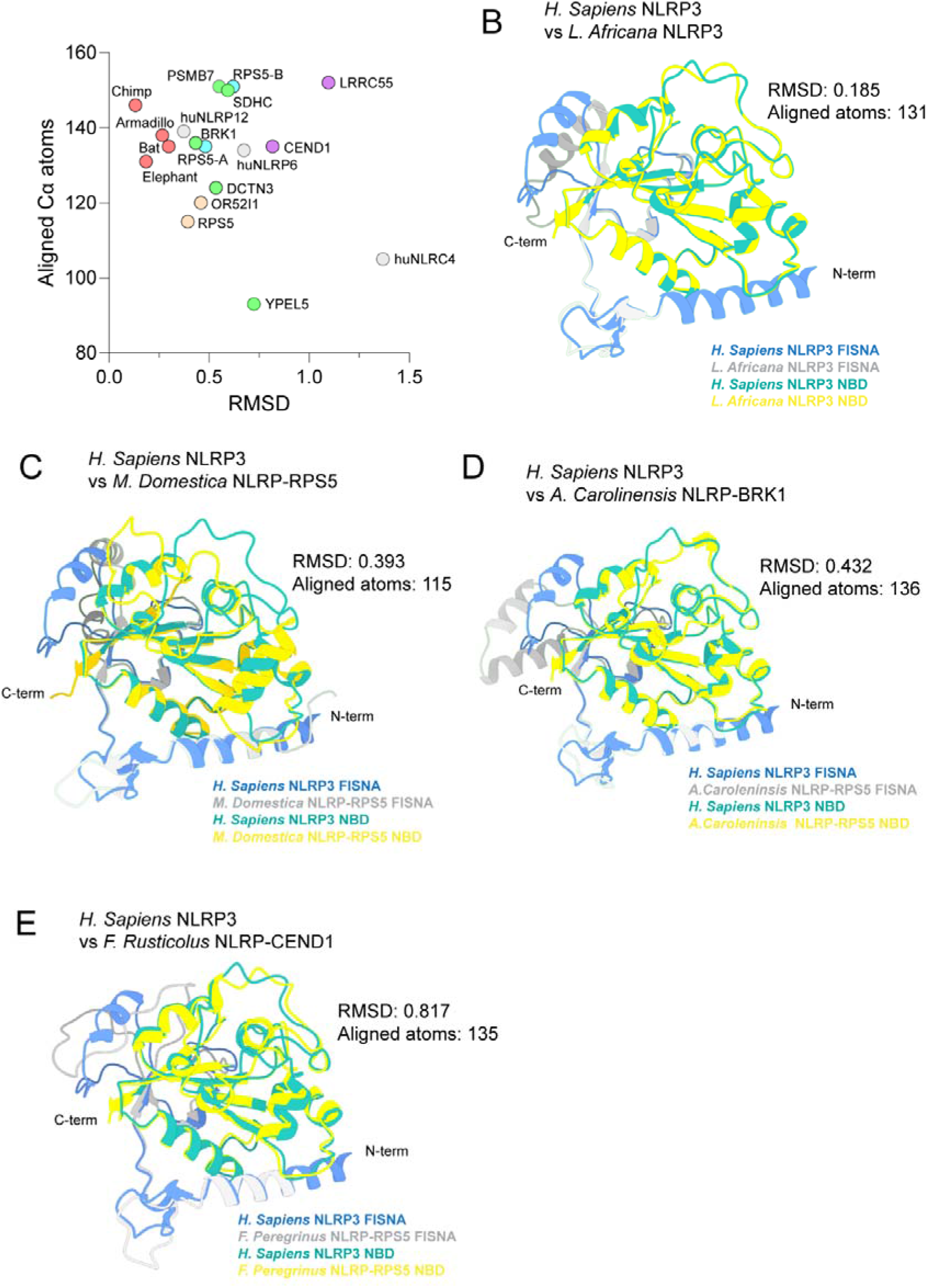
Structural alignments of the FISNA-NBD region between human NLRP3 and non-eutherian NLRP proteins. **A)** Graph plotting RMSD values against the number of aligned Cα atoms for each eutherian or non-eutherian NLRP protein aligned against human NLRP3. Data points are colour coded based on vertebrate lineage. Alignments were performed in PyMol, with visualisations of the aligned regions generated in ChimeraX. Graphics of alignments between *H. Sapiens* NLRP3^154-370^ and **B)** *L. Africana* NLRP3^147-372^ **C)** *M. Domestica* NLRP-RPS5^134-359^ **D)** *A. Carolinensis* NLRP-BRK1^110-339^ or E) *F. Peregrinus* NLRP-CEND1^133-359^. RMSD values and the number of aligned Cα atoms are indicated on each panel.

**Supplementary Figure 7.**
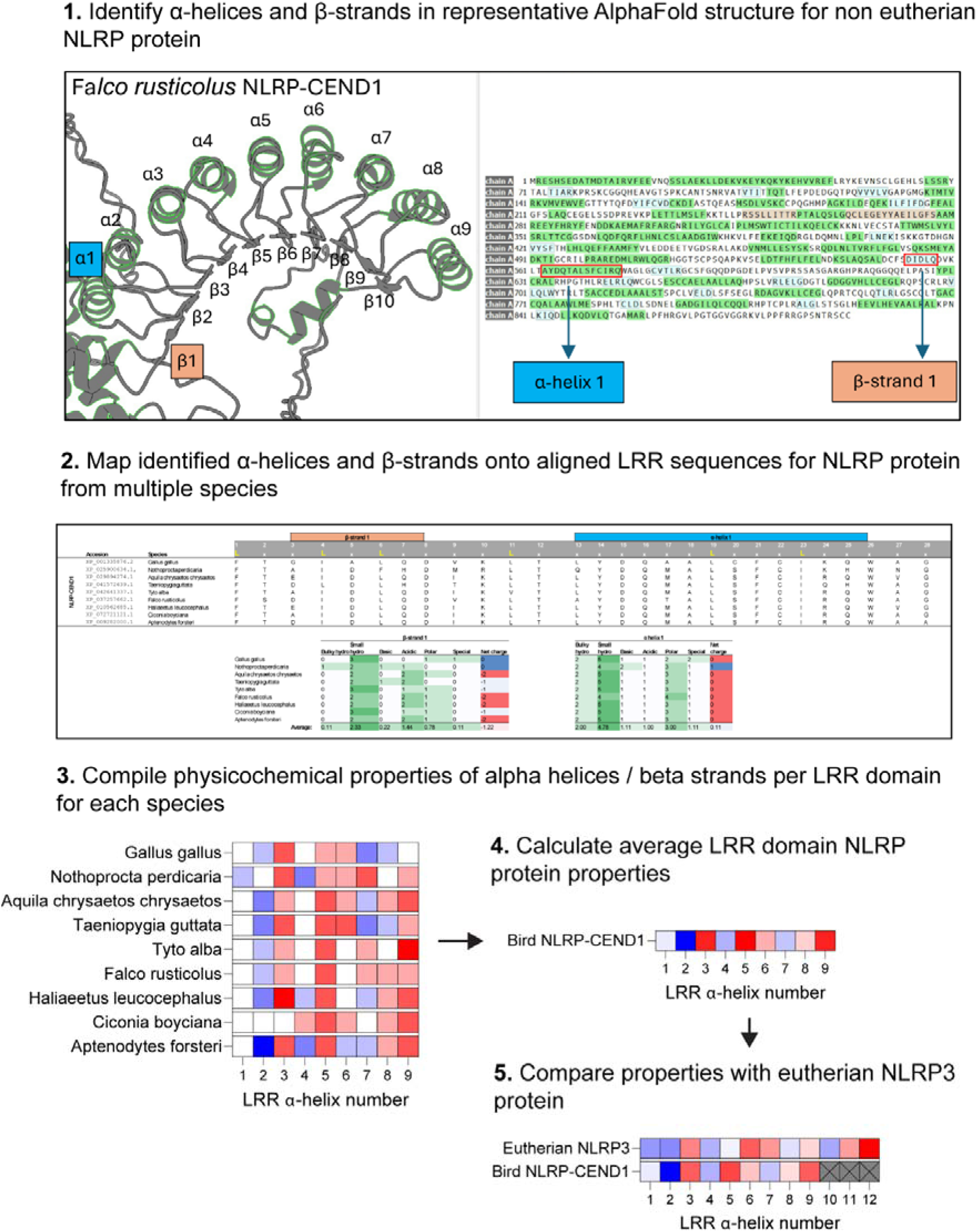
Overview of the workflow used to analyse the physicochemical properties of NLRP LRR domains.

**Supplementary Figure 8.**
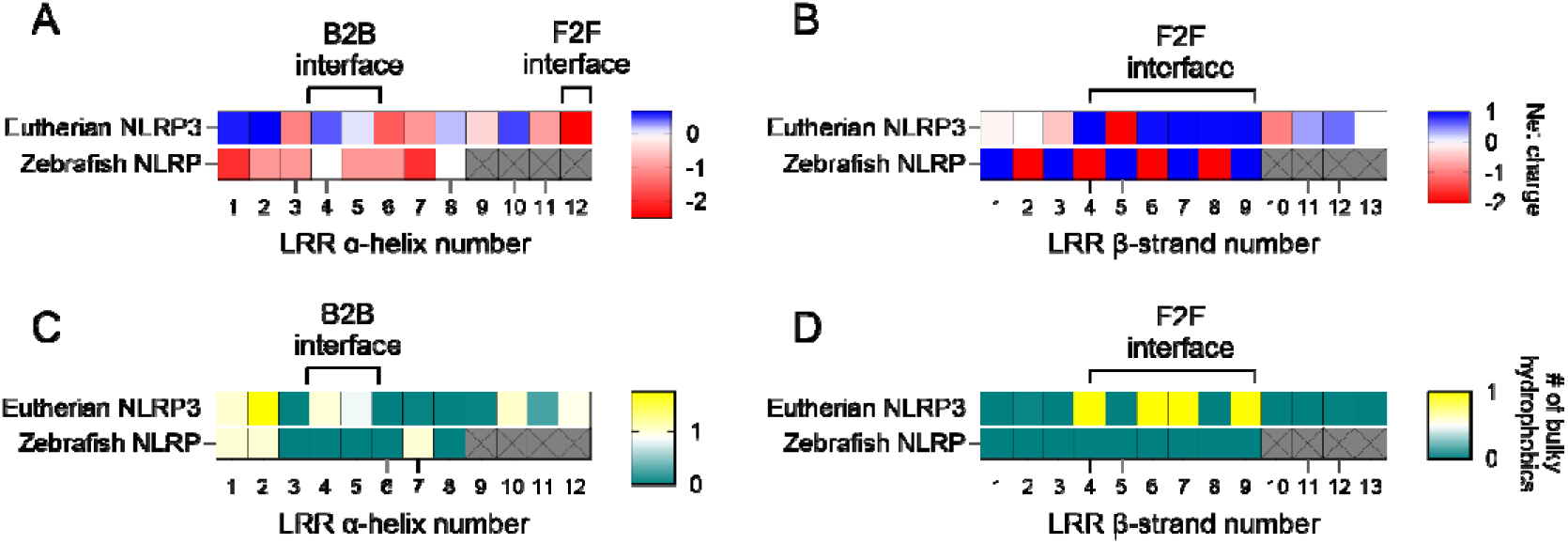
LRR domain analysis demonstrates the absence of conserved F2F and B2B interfaces in an experimentally validated zebrafish NLRP3-like protein (QEJ65894.1). Analysis of the net charge on each LRR **A)** alpha helix or **B)** beta strand for eutherian NLRP3 or zebrafish NLRP3-L. Analysis of the number of bulky hydrophobic residues on each LRR **C)** alpha helix or **D)** beta strand.

**Supplementary Figure 9.**
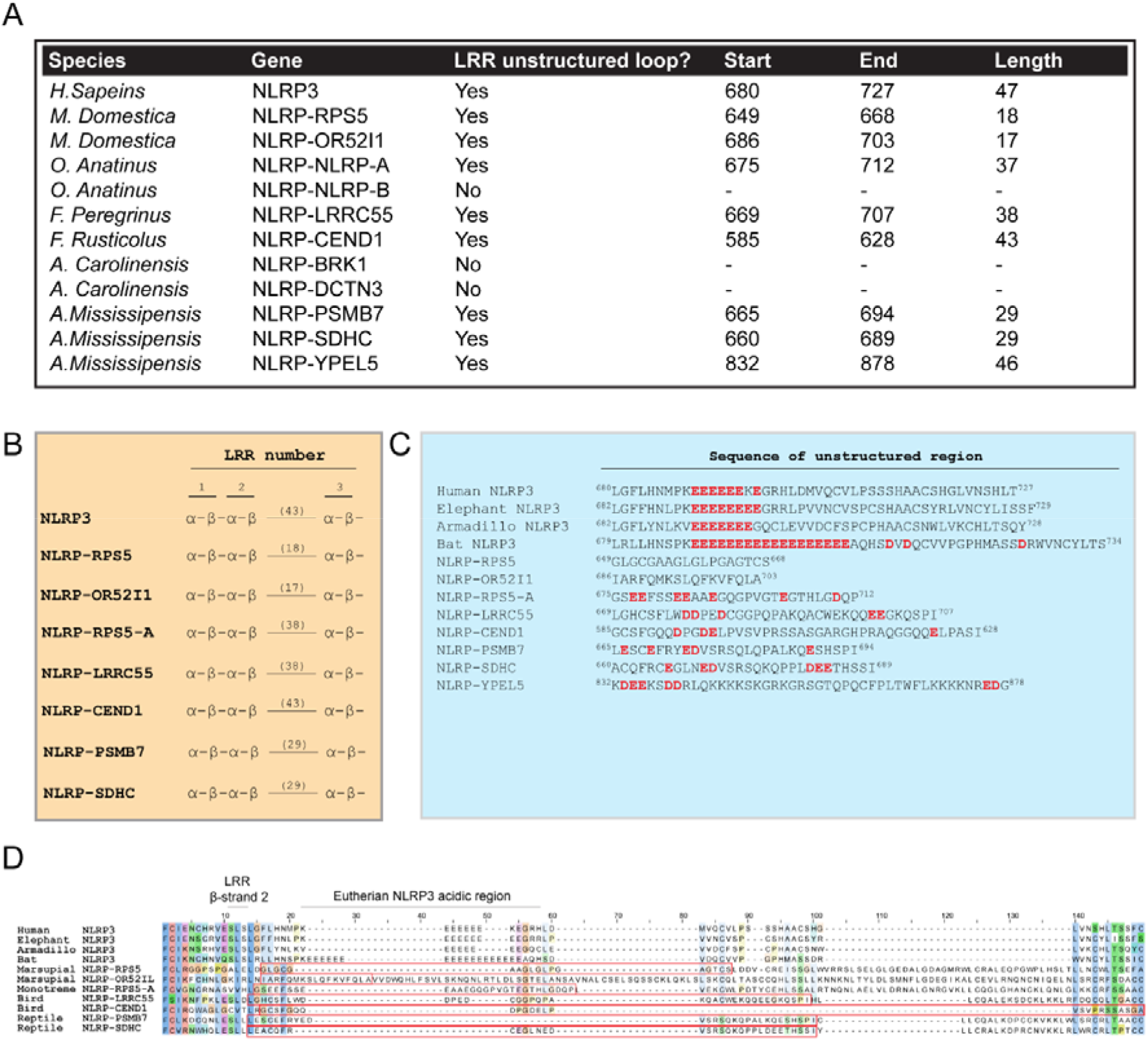
Non-eutherian NLRP proteins lack a region comparable to the eutherian NLRP3 acidic loop. **A)** Table detailing the presence or absence of an unstructured loop within the LRR domains of the indicated NLRP proteins alongside the position and sequence of unstructured loops if present. **B)** Table showing the positions of unstructured loops in relation to each α-helix and β-strand within the LRR domain. Unstructured loops are indicated by an extended solid line with length in amino acids annotated above. Except for NLRP-YPEL5 (not shown), all identified loops begin at the transition between the 2^nd^ LRR beta strand and 3^rd^ LRR alpha helix. **C)** Sequences of unstructured loops present in eutherian NLRP3 and non-eutherian NLRP proteins with acidic residues highlighted in red. A minimum of six consecutive acidic residues are found in eutherian NLRP3 proteins which contrasts with a maximum of three found in non-eutherian proteins. **D)** Sequence alignments of the regions containing the identified unstructured loops (boxed in red).

**Supplementary Figure 10.**
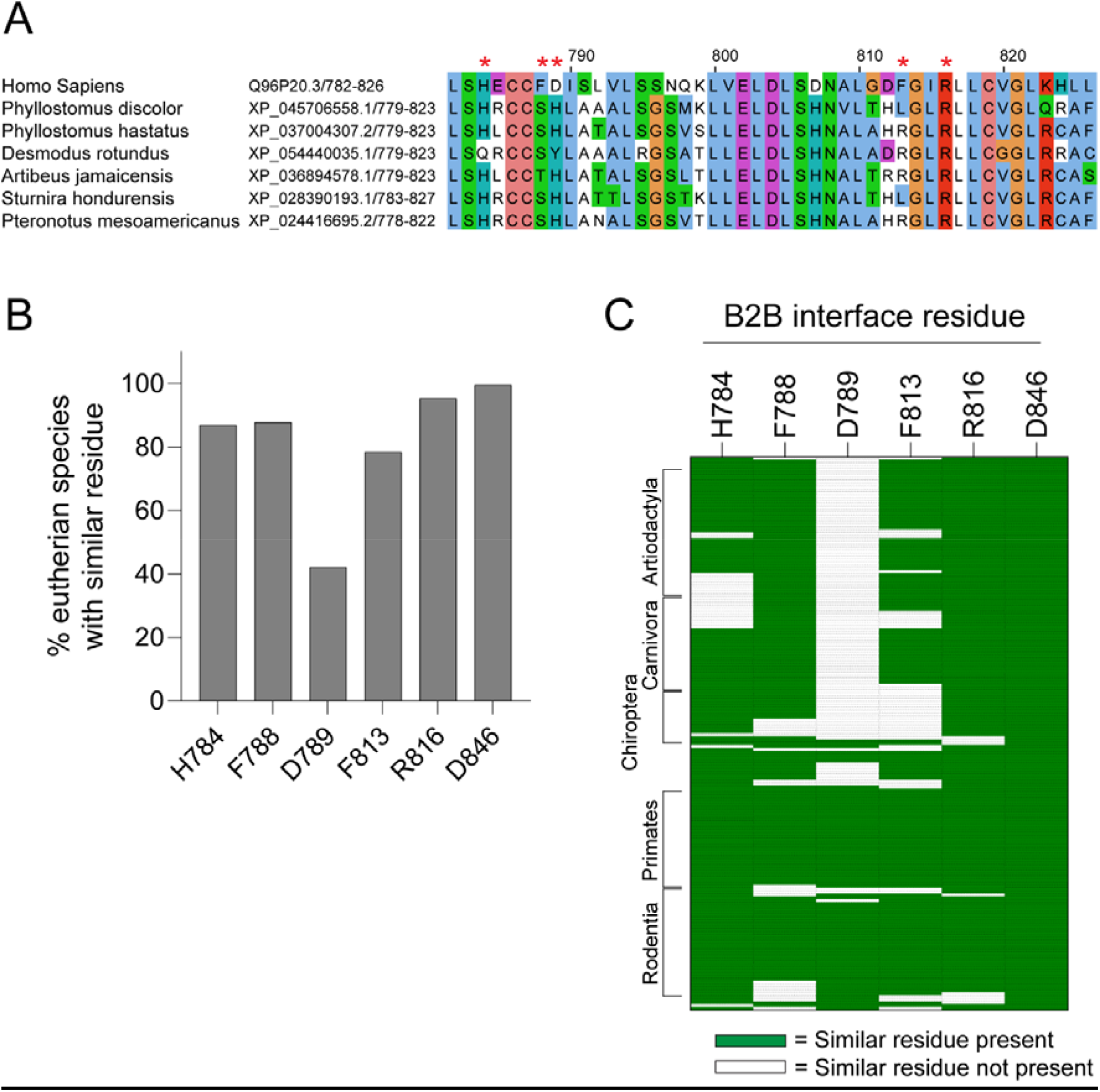
Sequence analysis of residues involved in formation of the back-to-back NLRP3 interface. **A)** Sequence alignments of the back-to-back region in bats lacking conservation of interface residues. Red asterisks indicate amino acids identified in cryo-EM structures of NLRP3 that participate in formation of the back-to-back interface. **B)** Bar chart showing the percentage of eutherian NLRP3 species containing a similar back-to-back interface residue found in the human NLRP3 sequence at the indicated position. **C)** Heat map showing residue similarity at the indicated amino acid position per eutherian species containing NLRP3. The presence of a similar residue is coloured green, the absence of a similar residue is coloured white.

## Table legends

**Supplementary Table 1.** Sequence identifiers for the eutherian NLRP3 and non-eutherian NLRP proteins analysed.

**Supplementary Table 2.** Sequence identifiers for amphibian and fish NLR proteins.

**Supplementary Table 3.** Sequence alignments of eutherian NLRP3, marsupial, monotreme, reptile, bird NLRPs and eutherian NLRP12.

